# Speech recognition from MEG data using covariance filters

**DOI:** 10.1101/2023.06.22.546174

**Authors:** Vitaly Verkhlyutov, Victor Vvedensky, Konstantin Gurtovoy, Evgenii Burlakov, Olga Martynova

## Abstract

Speech recognition from EEG and MEG data is the first step in the development of BCI and AI systems for further use in the decoding of inner speech. Great achievements in this direction have been made with the use of ECoG and stereo-EEG. At the same time, there are few works on this topic on the analysis of data obtained by noninvasive methods of brain activity registration. Our approach is based on the evaluation of connections in the sensor space with the extraction of the MEG connectivity pattern specific to a given segment of speech. We tested our method on 7 subjects. In all cases, our processing pipeline was sufficiently reliable and worked either without recognition errors or with few errors. After ”training” the algorithm is able to recognize a fragment of spoken speech in a single presentation. For recognition, we used MEG recording segments of 50-1200 ms from the beginning of the word. A segment of at least 600 ms was required for high-quality recognition. Intervals longer than 1200 ms degraded the quality of recognition. Band-pass filtering of MEG showed that the quality of recognition is higher when using the gamma frequency range compared to the low-frequency range of the analyzed signal.

## 1 Introduction

Decoding of inner speech and speech stimuli from brain activity data is an urgent task for theoretical and applied purposes of modern neurophysiology. Within this direction, researchers are trying to solve the problem of compensation for lost functions in various types of disorders of speech reproduction and perception at the cortical level, which has direct relevance to BCI. At the same time, the study of this issue helps to move towards the improvement of AI systems. Significant progress has been made with intracranial ECoG recordings [1] and stereo EEG [2]. However, invasive techniques have a limited range of applications. Recent studies have shown that decoding macroscopic fMRI data using a trained language model can quite accurately decode internal speech based on semantic information [3].

Other non-invasive recording methods, such as EEG and MEG, have proven that speech perception and playback affect rhythmic [4], [5], [6] and evoked [7] brain electrical activity. Thus, there are all prerequisites for speech decoding based on MEG and EEG data. However, for the analysis of brain activity in this case [8] neural network technologies are used, the results of which are difficult to interpret. For these purposes, we propose to use a simpler technique to investigate the connectivity of MEG in the sensor space, which is based on observations showing a remarkable similarity of the current MEG activity on lead clusters when listening to words, as well as the dynamic reorganization of these clusters when recognizing the semantic meaning of a speech stimulus [4].

## 2 Measurements

### Subjects

Seven volunteer subjects (four men and three women) participated in the pilot study aimed at testing the technique. One of the subjects was left-handed at the age of 23. The average age of the young right-handed subjects was 23.8 ±0.5 years. The elderly right-handed subject was 67 years old. All subjects had no history of neurological or psychiatric disorders. The study was carried out in accordance with The Code of Ethics of the World Medical Association (Declaration of Helsinki) for experiments involving humans, and the Ethical committee of the Institute of Higher Nervous Activity and Neurophysiology of the RAS approved the research protocols (Protocol No. 5 of December 2, 2020). The studies took place from 12 to 15 o’clock.

### Stimuli

The subject was presented with three series of speech stimuli in the form of Russian adjectives. Each series included eight original words, which were repeated five times. All forty words were randomly mixed. Before each series, three words from the same series of words were presented to adapt the subject, but the recorded data from these presentations were not considered for analysis. The series of words differed in sound duration and were 600, 800, and 900 ms, respectively. The loudness of the sound was selected for each subject and ranged from 40 to 50 dB. The frequency of the digitized words as an audio file did not exceed 22 kHz. After a word was presented, the subject had to press the hand-held manipulator button if he understood the meaning of the presented word. Pressing the button after 500 ±100 ms (randomized) was followed by the next stimulus, but no later than 2000 ms after the previous presentation.

### Experimental procedure

Before the experiment, the coordinates of the anatomical reference points (left and right preauricular points and the bridge of the nose) were determined using the FASTRAK 3D digitizer (Polhemus, USA), as well as indicator inductance coils attached to the subject’s scalp surface in the upper part of the forehead and behind the auricles. During the experiment, the subject was in a magnetically shielded multilayer permalloy chamber (AK3b, Vacuumschmelze GmbH, Germany) and his head was placed in a fiberglass helmet, which is part of a fiberglass Dewar vessel with a sensor array immersed in liquid helium. The test subject was seated so that the surface of the head was as close as possible to the sensors. To avoid artifacts, sound stimuli were delivered through a pneumatic system delivering sound from a standard audio stimulator. The stimulator was programmed using Presentation software (USA, Neurobehavioral Systems, Inc). The subject was asked to relax and close his eyes. His right hand touched a console with buttons. He had to press one key with his index finger after recognizing the heard word. At the end of the series of presentations, the subject was allowed to rest for 1-2 minutes.

### Registration

Registration parameters are described on the Zenodo service. The data (MEG and MRI) are also available there [9].

## 3 Data analysis

We did not use any additional signal processing methods except MaxFilter and bandpass filtering to determine the contribution of the delta-gamma of the MEG frequency range to the correctness of word recognition. Actual segments of the MEG were identified using the marks of the beginning of the sounding of words. These segments were used to build covariance matrices, calculating the Pearson correlation of each registration channel with each and thus forming a covariance vector for the original word

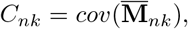

where 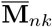 is the vector of the MEG data for the *k*-th repeat of the *n*-th word (*n* = 1, …, *N, k* = 1, …, *K*). The covariation matrices were averaged for all words by subtracting from them the averaged matrix for each word with replacing the principal diagonal elements by zeros and setting to zero all elemets that less than 0.7 in the resulting matrices:

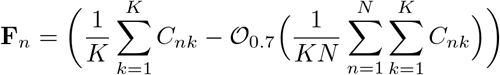

where the operator 𝒪_*x*_ replaces the principal diagonal elements by zeros and setting to zero all elemets that less than *x*. The resulting patterns were used to calculate the weights of newly presented words. The weight *w* of a word with the covariation matrix *C* can be assessed with respect to the 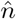 -th word filter **F**_*n*_ as

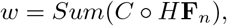

where the functional *Sum* translates any matrix to the sum of its elements, the binary operation ◦ represents the elementwise product of two matrices (of the same dimension), and *H* is the elementwise Heaviside function. If the weight of a recognised word exceeded the weight of all others, then the word was considered recognised. Thus, the number of the recognised word can be found from the relation

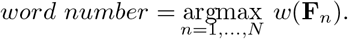

Weights were calculated for all 40 words. The 5 maximum weight values belonged to the recognized word. The recognition error was considered to be weight reduction below the maximum weight for all presentations of all the remaining 35 other words. Thus, the system could make a maximum of 5 errors when recognizing one word and 40 errors when recognizing 8 original words. At the same time, we could evaluate the success rate of recognition as a percentage. At 100% recognition, all 5 identical words were identified in a sequence of 40 words. One error reduced the recognition success score by 2.5%. The described algorithm is implemented by Matlab [10].

## 4 Results

Correlation analysis of the one-second segments of MEG showed the range of the correlation coefficients from *r <* 0.9 to *r >−* 0.9 (Fig. 1). However, we only used *r* values that are grearter than 0.7 for the analysis.

**Fig. 1.**
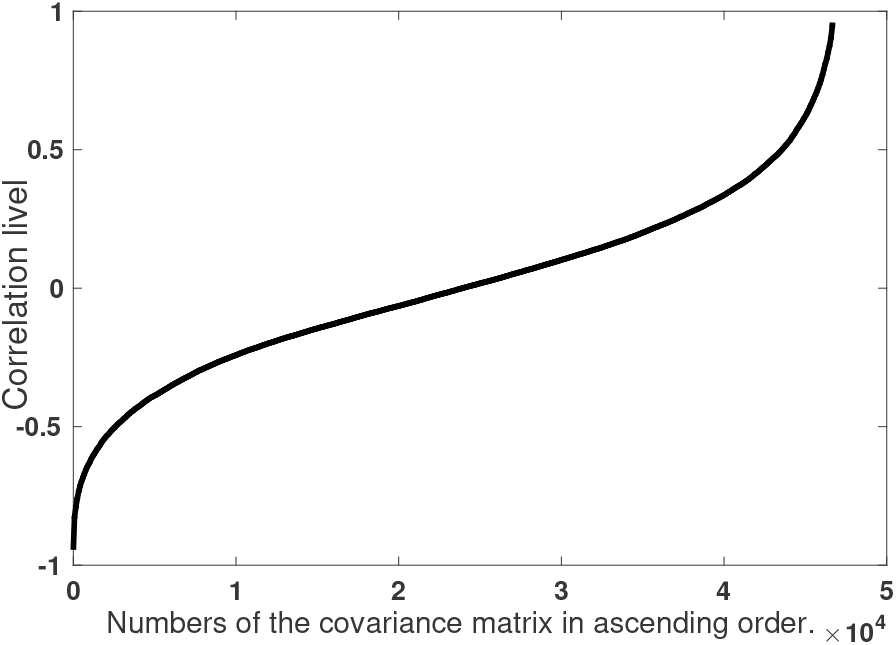
Correlation coefficients when comparing each sensor to each from a one-second segment of MEG at the time of word sound. Mirror elements have been removed.

Weights were obtained for all 24 words in three series (Fig. 2) for each subject. It was not always possible to successfully recognize a target word with a weight close to or less than one of the background words or words of choice. In this case, recognition was considered inaccurate. An attempt to select the optimal length of the MEG segment for recognition was made (see Fig. 3).

**Fig. 2.**
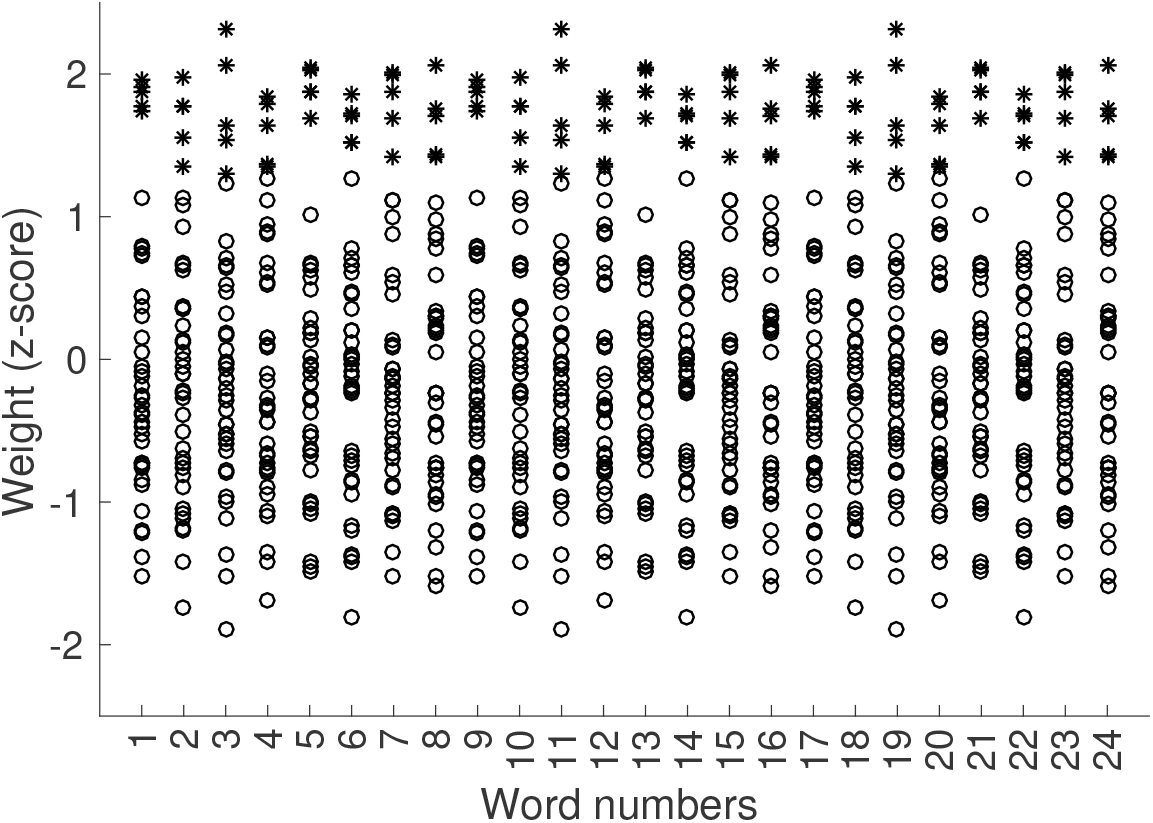
Normalized weights for 24 original words (5 presentations for each word) for 3 series of presentations of subject V4. Each column contains 40 weights. Asterisks show 5 recognized words, circles show background words. If there are less than 5 stars, then there is layering, and the weights have very close values.

**Fig. 3.**
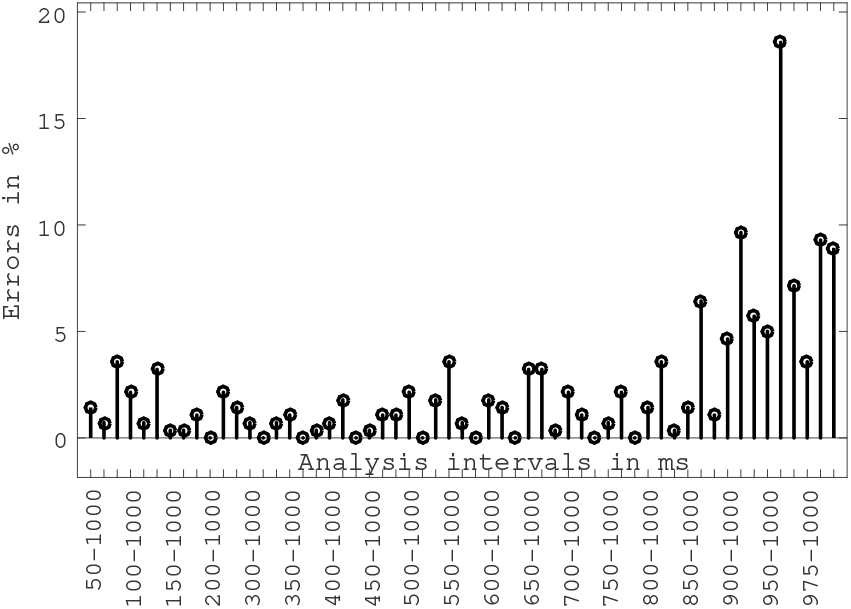
The effect of misrecognition, when the analyzed MEG segment is reduced from 50-1000 ms to 975-1000 ms from the beginning of the word in the subject V1. The unit stem denotes the percentage of errors in the recognition of one set of 8 original words. Intervals are indicated only for the first set of three.

Reducing the analyzed MEG segment had little effect on the quality of recognition until the segment of 850-1000 ms. Fig. 3 shows the percentage of errors for subject V1 for three series of presentations separately. At intervals from 200-1000 ms to 750-1000 ms, a zero error rate is observed in the individual series (the circle lies on the zero line). By increasing the same analyzed MEG interval from 0-50 ms to 0-1200 ms, the quality of recognition began to increase with durations from 0 to 600 ms (Fig.4). Allocating 100 ms segments for analysis caused a deterioration of word recognition quality, especially at segments from 200 to 300 ms and from 1000 to 1100 ms (Fig. 5).

**Fig. 4.**
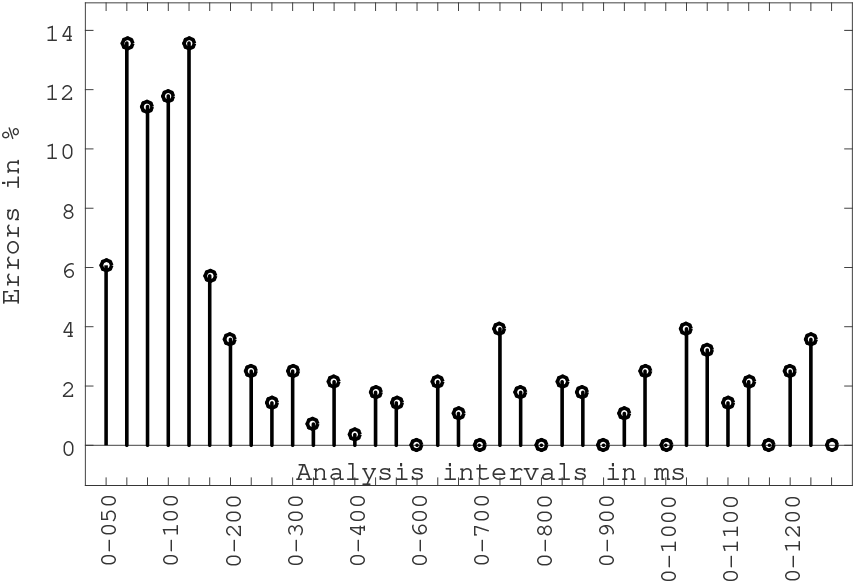
The effect of misrecognition, when the analyzed MEG segment is increased from 0-50 ms to 0-1200 ms from the beginning of the word in subject V1. The unit stem denotes the percentage of errors in the recognition of one set of 8 original words. Intervals are indicated only for the first set of three.

**Fig. 5.**
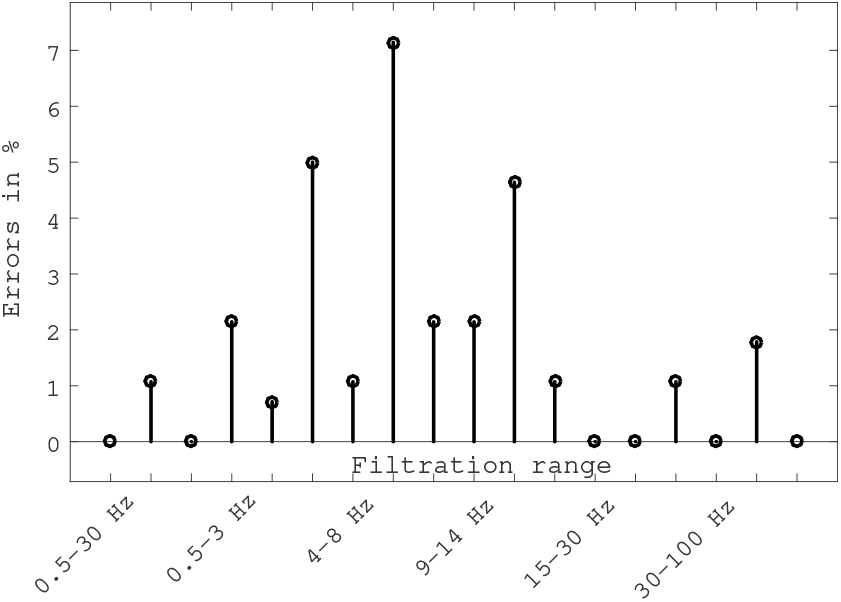
The effect of misrecognition, when bandpass filtering the interval 200-1000 ms from the beginning of the word in the subject V1 in the ranges 0.5-30 Hz, 0.5-3 Hz, 4-8 Hz, 9-14 Hz, 15-30 Hz, 30-100 Hz. The single stem denotes the percentage of errors in the recognition of one set of 8 original words. Intervals are marked only for the first set of three.

We evaluated the quality of recognition in all subjects (Table 1). We did not find any tendency depending on age, gender, and dominant hand (subject V3 was elderly and subject V5 was left-handed).

**Table 1.**
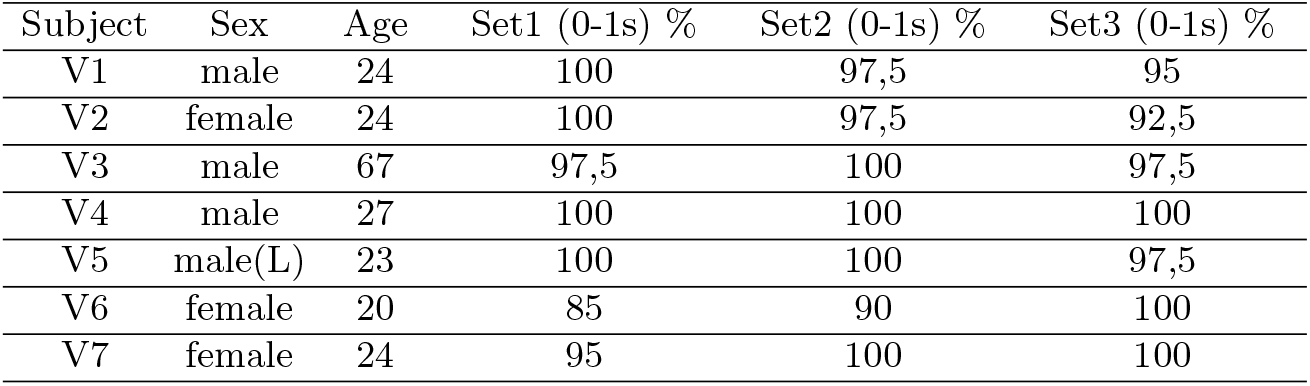
Percentage of correct recognition in 7 subjects using a 0-1000 ms interval of unfiltered MEG from word onset for three sets of words.

| Subject | Sex | Age | Set1 (0-1s) % | Set2 (0-1s) % | Set3 (0-1s) % |
| --- | --- | --- | --- | --- | --- |
| V1 | male | 24 | 100 | 97,5 | 95 |
| V2 | female | 24 | 100 | 97,5 | 92,5 |
| V3 | male | 67 | 97,5 | 100 | 97,5 |
| V4 | male | 27 | 100 | 100 | 100 |
| V5 | male(L) | 23 | 100 | 100 | 97,5 |
| V6 | female | 20 | 85 | 90 | 100 |
| V7 | female | 24 | 95 | 100 | 100 |

## 5 Discussion

An important factor for the behavior of neuronal populations is their synchro-nization, which allows many neurons to work in parallel and process many properties of the input signal simultaneously, establishing their multiple connections with other mental objects and their properties [11]. In our experiments, we observe that some populations are active in the perception of any word, and some are specific to a particular word. A speech recognition system can be based on the specificity property. In doing so, we investigate only amplitude connectivity, which is due to distant connections [12] as opposed to phase connectivity, which in turn is provided by local interactions. The presence of phase synchronization in our experiments proves the presence of both positively and negatively correlated data. Phase coupling indicates possible effects associated with the rotation of current dipoles, which are due to cortical traveling waves [13]. The effectiveness of individual segments for decoding is possibly related to implicit perception, which allows the brain to view part of a phrase as having complete meaning [14]. Bandpass filtering has shown that high-frequency components are most effective for decoding, though along with other frequency bands, due to long-range interaction (through myelinated fibers) between local electrical brain sources [15].

## References

1. Anumanchipalli, G.K., Chartier, J., Chang, E.F.: Speech synthesis from neural decoding of spoken sentences. Nature, 568(7753), 493–498 (2019). https://doi.org/10.1038/s41586-019-1119-1

2. Norman-Haignere, S.V., Long, L.K., Devinsky, O., Doyle, W., Irobunda, I., Merricks, E.M., Mesgarani, N.: Multiscale temporal integration organizes hierarchical computation in human auditory cortex. Nature Human Behaviour, 6(3), 455–469 (2022). https://doi.org/10.1038/s41562-021-01261-y

3. Tang, J., LeBel, A., Jain, S., Huth, A.G.: Semantic reconstruction of continuous language from non-invasive brain recordings. Nature Neuroscience. (2023). https://doi.org/10.1038/s41593-023-01304-9

4. Vvedensky, V., Filatov, I., Gurtovoy, K., Sokolov, M.: Alpha Rhythm Dynamics During Spoken Word Recognition. Studies in Computational Intelligence, 1064, 65–70 (2023). http://doi.org.10.1007/978-3-031-19032-27

5. Lizarazu, M., Carreiras, M., Molinaro, N. (2023). Theta-gamma phase-amplitude coupling in auditory cortex is modulated by language proficiency. Human Brain Mapping, 44(7), 2862–2872. https://doi.org/10.1002/hbm.26250

6. Neymotin, S.A., Tal, I., Barczak, A., O’Connell, M.N., McGinnis, T., Markowitz, N., Lakatos, P.: Detecting Spontaneous Neural Oscillation Events in Primate Auditory Cortex. Eneuro, 9(4), ENEURO.0281-21 (2022). https://doi.org/10.1523/ENEURO.0281-21.2022

7. Anurova, I., Vetchinnikova, S., Dobrego, A., Williams, N., Mikusova, N., Suni, A., Palva, S.: Event-related responses reflect chunk boundaries in natural speech. NeuroImage, 255, 119203 (2022). https://doi.org/10.1016/j.neuroimage.2022.119203

8. Dash, D., Ferrari, P., Wang, J.: Decoding Imagined and Spoken Phrases From Non-invasive Neural (MEG) Signals. Frontiers in Neuroscience, 14, 290 (2020). https://doi.org/10.3389/fnins.2020.00290

9. Verkhlyutov, V.: MEG data during the presentation of Gabor patterns and word sets. ZENODO, 7458233 (2022). https://zenodo.org/record/7458233

10. https://github.com/BrainTravelingWaves/22SpeechRecognition

11. Defossez, A., Caucheteux, C., Rapin, J., Kabeli, O., King, J.-R. Decoding speech from non-invasive brain recordings. ArXiv, 2208.12266, 1–15. (2022). http://arxiv.org/abs/2208.12266

12. Rolls, E.T., Deco, G., Huang, C.-C., Feng, J.: The human language effective connectome. NeuroImage, 258, 119352 (2022). https://doi.org/https://doi.org/10.1016/j.neuroimage.2022.119352

13. Sato, N.: Cortical traveling waves reflect state-dependent hierarchical sequencing of local regions in the human connectome network. Scientific Reports, 12(1), 334 (2022). https://doi.org/10.1038/s41598-021-04169-9

14. Liaukovich, K., Ukraintseva, Y., Martynova, O.: Implicit auditory perception of local and global irregularities in passive listening condition. Neuropsychologia, 165(July 2020), 108129 (2022). https://doi.org/10.1016/j.neuropsychologia.2021.108129

15. Proix, T., Delgado Saa, J., Christen, A., Martin, S., Pasley, B.N., Knight, R.T., Giraud, A.-L.: Imagined speech can be decoded from low- and cross-frequency intracranial EEG features. Nature Communications, 13(1), 48 (2022). https://doi.org/10.1038/s41467-021-27725-3

